# DisPhaseDB, an integrative database of diseases related variations in liquid-liquid phase separation proteins

**DOI:** 10.1101/2022.02.03.479026

**Authors:** Alvaro Navarro, Fernando Orti, Elizabeth Martínez-Pérez, Franco Simonetti, Javier Iserte, Cristina Marino-Buslje

## Abstract

**Motivation:** Proteins involved in liquid-liquid phase separation (LLPS) and membraneless organelles (MLOs) are recognized to be decisive for many biological processes and also responsible for several diseases. The recent explosion of research in the area still lacks tools for the analysis and data integration among different repositories. Currently, there is not a comprehensive and dedicated database that collects all disease-related variations in combination with the protein location, biological role in the MLO and all the metadata available for each protein and disease. Disease related protein variants and additional features are dispersed and the user has to navigate many databases, with different focus, formats and often not user friendly.

**Results:** We present DisPhaseDB, a database dedicated to disease related variants of LLPS proteins and/or are involved in MLOs. It integrates 10 databases, contains 5.741 proteins, 1.660.059 variants and 4.051 disease terms. It also offers intuitive navigation and an informative display. It constitutes a pivotal starting point for further analysis, encouraging the development of new computational tools.

**Availability and Implementation:** The database is freely available at http://disphasedb.leloir.org.ar.

**Graphical abstract:** 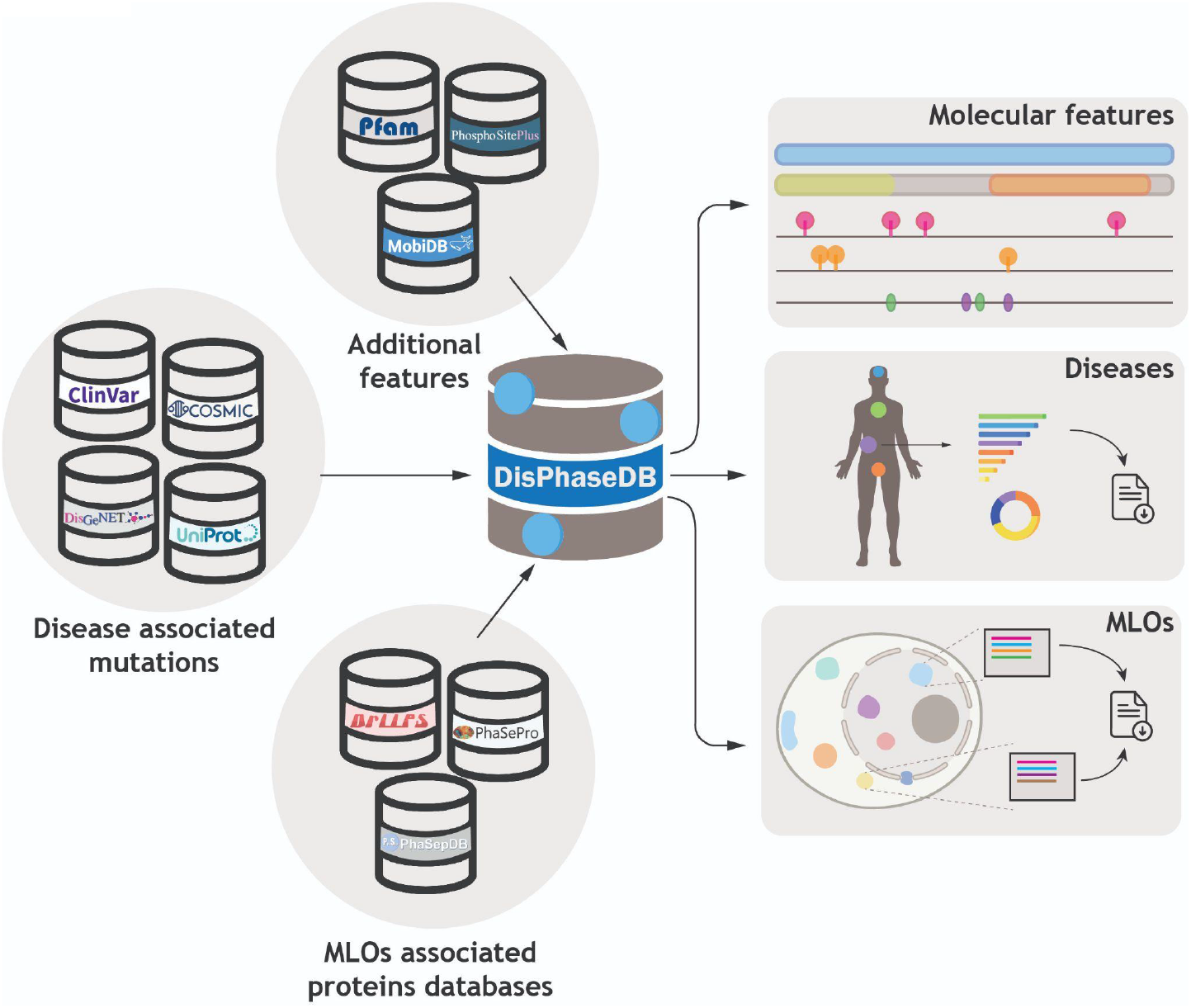

## Introduction

Cells compartmentalize biological processes to achieve spatial and temporal control over biochemical reactions. This is accomplished through both membrane bound and membraneless organelles (MLOs). MLOs are formed through the process of Liquid-liquid phase separation (LLPS) in which a liquid demixes in two phases where one phase is enriched in particular macromolecules, while depleted in others (Mao, Zhang and Spector, 2011; Banani *et al*., 2017; Sanders *et al*., 2020) Examples are Nucleolus, Cajal bodies, and nuclear speckles in the nucleus and stress granules, P granules and P-bodies in the cytoplasm (Mao, Zhang and Spector, 2011; Banani *et al*., 2017), among others. These structures play diverse roles in various biological processes such as organization of the cytoplasm and nucleoplasm, regulation of gene expression, signaling, transport and compartmentalization (Gomes and Shorter, 2019; Su, Mehta and Zhang, 2021). However, they are also increasingly implicated in several complex human diseases (Shin and Brangwynne, 2017; Lenard *et al*., 2021; Su, Mehta and Zhang, 2021). Examples of abnormal LLPS have been implicated in cancer, neurodegenerative and infectious diseases among others (Harrison and Shorter, 2017; Alberti and Dormann, 2019; Prasad *et al*., 2019; Ryan and Fawzi, 2019; Tsang *et al*., 2020). Therefore, it is not surprising that a perturbation in proteins that undergo LLPS, like a single nucleotide variant (SNV), gene copy number variation (CNV), protein mutation and post-translational modifications (PTMs) can upset the fine tuned process of MLO formation, stability and dynamics (Murakami *et al*., 2015; Gopal *et al*., 2017; Tang, 2017; Hofweber and Dormann, 2019; Wu *et al*., 2020; Lenard *et al*., 2021; Luo *et al*., 2021; Schisa and Elaswad, 2021; Specht, 2021).

Proteins that undergo LLPS are often intrinsically disordered or have disordered regions (IDPs and IDRs, respectively), they might also have a biased amino acid composition or low-complexity regions (LCRs), and are therefore highly dynamic (van der Lee *et al*., 2014; Boeynaems *et al*., 2018; Luo, Wu and Li, 2021). To a large extent, these regions are responsible for the separation in phases, although other types of regions or domains can also be found in proteins that separate into phases (Vernon *et al*., 2018; Dignon, Best and Mittal, 2020; Luo, Wu and Li, 2021). There are several molecular interaction types contributing to LLPS, such as multivalent protein-protein and protein-DNA/RNA interactions, dinamically transcient interacting regions as IDRs, LCR and prion-like, aggregation, coacervation, electrostatic, cation-π and π-π interactions, among others (Mittag and Parker, 2018). Mapping mutations to structural features could help to understand mechanisms involved in the formation of pathological aggregates. As an example, it was shown that mutations in the prion-like domains (PLDs) of several proteins are involved in neurodegenerative diseases such as amyotrophic lateral sclerosis (ALS), frontotemporal dementia (FTD), and multisystem proteinopathy (Harrison and Shorter, 2017).

Numerous experimental methods have been developed or repurposed to study the LLPS process and proteins involved, such as fluorescence recovery after photobleaching (FRAP), nuclear magnetic resonance spectroscopy (NMR), immunofluorescence, fluorescence correlation spectroscopy (FCS), and many others (Gadd *et al*., 2008; Brangwynne *et al*., 2009; Schmidt and Görlich, 2015; Alderson and Kay, 2021). However, there are still not enough bioinformatics tools and databases to study them, much less in the context of human diseases. We hypothesize that this is in part due to the lack of centralized data repositories, the low agreement among existing ones, the scarcity of dedicated cross-referenced databases and, the poor scalability offered for large integrative analysis of phase separating proteins.

It was shown that the agreement between 4 dedicated databases of LLPS proteins (Li *et al*., 2019; Mészáros *et al*., 2020; Ning *et al*., 2020; You *et al*., 2020) is rather poor, sharing 42 human proteins out of 4,367, proving that none of the four databases taken alone provides enough data to enable a meaningful analysis (Orti *et al*., 2021), added to the fact that they do not focus on protein variations in diseases.

To cover this gap, we present **DisPhaseDB**, an integrative database focused on disease variations in LLPS proteins. The database encompasses all known phase separating proteins, including Drivers, Clients, Regulators and other MLOs experimentally associated proteins together with their disease related variations. We expect our database to be of interest for researchers studying MLOs, LLPS proteins, diseases, proteins for targeting therapies, specific MLO components in a disease and also for computational groups developing methods to understand sequence-function relationships and mutational impact.

## Methods

### Selection of proteins involved in LLPS and MLO associated

Our starting point was an integrated set of MLO associated proteins that were collected in a previous group effort (Orti *et al*., 2021). It consists of the entries of four databases of LLPS and MLOs associated proteins that were compiled, merged, completed and stored in a local database: **PhaSePro** (Mészáros *et al*., 2020), **PhaSepDB** (You *et al*., 2020) **DrLLPS** (Ning *et al*., 2020) **and LLPSDB** (Li *et al*., 2019). This set is periodically updated with the databases’ new releases. The consolidated dataset is available at http://mlos.leloir.org.ar/ (Orti *et al*., 2021).

The role of the proteins in the LLPS process and their association with the MLOs, is taken from the annotation of the source database. There are four types of Protein roles: Drivers, Regulators, Clients and Unassigned when no database describes its role. In addition, we grouped their experimental evidence supporting the roles as low throughput and high throughput for user evaluation of their confidence.

### Mutation collection

We obtained human coding variants from **ClinVar** release 20200402 (Landrum *et al*., 2018), a public archive of human genetic variants and their interpretation with respect to a clinical condition or phenotypes, along with supporting evidence for such association. **DisGeNET** (Piñero *et al*., 2020) offers several datasets based on gene-disease associations (GDAs) and variant-disease associations (VDAs). For our database we took mutations from the curated VDA dataset (October 2020), which at the same time integrates variants from UniProt, ClinVar, GWASdb (Li *et al*., 2012) and GWAS catalog (Buniello *et al*., 2019).

From **UniProt** (UniProt Consortium, 2021) we used the dataset of human variants that are manually annotated in UniProtKB/Swiss-Prot (release-2021_02). Lastly, **COSMIC** release v94 was used to obtain the coding point mutations in human cancers (Tate *et al*., 2019).

In all cases, we mapped variants with genomic coordinates from the human genome assembly GRCh38 onto the canonical protein sequence. Disease and other altered phenotypic effects annotations in ClinVar, COSMIC, DisGeNET and UniProt are not consistent between databases nor within the same database. They are frequently cross referenced to one or many ontologies that collect medical terms, and/or diseases, such as Disease Ontology (DO) (Schriml *et al*., 2019), the Human Phenotype Ontology (HPO) (Köhler *et al*., 2021), Medical Subject Headings (MeSH) (Nelson, 2009), Medical Genetics (MedGen, https://www.ncbi.nlm.nih.gov/medgen/), The Monarch Merged Disease Ontology (MONDO) (Mungall *et al*., 2017), National Cancer Institute Thesaurus (NCI, https://ncim.nci.nih.gov/), Online Mendelian Inheritance in Man (OMIM) (Amberger *et al*., 2018), among others. In some cases there is no reference to any ontology. A mutation can be associated with several diseases and vice versa. In this context studying a variant, a protein or a disease is challenging. As an example, mutation R521C in FUS protein is associated with different diseases in different ontologies: Melanoma of skin (SNOMEDCT_US: 93655004), amyotrophic lateral sclerosis ALS6 (MEDCIN: 315716 and MedGen: C1842675) and Gastric Carcinoma (NCI: C4911). In addition, there are many synonyms annotated for the same disease in one ontology, as an example “Cancer of Stomach”, “Cancer of the Stomach”, “Carcinoma of Stomach”, “Gastric Cancer”, etc, are references to the same disease in NCI. Another case are synonyms in different ontologies, as example: Cutaneous Melanoma (MedGen: C0151779), Melanoma of skin (SNOMEDCT_US: 93655004) and “Melanoma, Cutaneous Malignant” (OMIM: 155600).

Finally, there are different grades of specificity of a disease that are referred to as different terms, as an example, “Acanthoma” is a type of “Skin Neoplasms”. Therefore, mapping all disease terminology into a single ontology is not feasible. So, to facilitate the user navigating through this tangle of terms in dozens of ontologies to study a variation or a protein, DisPhaseDB includes all available disease annotations and, when there are no references to an ontology, reference to the source mutation database.

### Additional information

We also included molecular features such as structural domains from Pfam database (Mistry et al., 2020), Intrinsically disordered Regions (IDRs) and Low-Complexity Regions (LCRs) from MobiDB (Piovesan *et al*., 2021), post-translational modifications (PTMs) retrieved from PhosphoSitePlus (Hornbeck *et al*., 2019) and Prion-like domains (PLDs) predicted by PLAAC (Alberti *et al*., 2009; Hornbeck *et al*., 2019). These features are displayed on the protein sequence using the “Feature-Viewer” tool to visualize positional data (Paladin *et al*., 2020).

### Implementation

The server backend consists of a http web-server developed in Python 3.8+ using the Flask framework and MySQL. The client web application was developed with the AngularJS framework.

## Results

### DisPhaseDB in numbers

We present DisPhaseDB, available at https://disphasedb.leloir.org.ar. DisPhaseDB contains 5.741 LLPS proteins, all of them with experimental evidence that supports their association to the MLOs. For these proteins we collected human disease mutations from up-to-date databases including **UniProt, ClinVar, DisGeNET** and **COSMIC**. After merging the four databases, the total number of unique coding variants (protein mutations) is 1.660.059. **COSMIC** contributes 1.464.124, **ClinVar** 221.097, **DisGeNET** 56.813 and **UniProt** 22.965. **Supplementary Figure 1** shows the overlap of the four protein variation resources, showing that all of them are needed to have a better landscape of mutation in LLPS proteins. The most common type are missense mutations, followed by synonyms mutations (66.57% and 23.41% respectively) **(Supplementary Figure 2)**.

On average, proteins in DisPhaseDB have around 200 mutations, although few proteins are exceptionally highly mutated **(Figure 1)**. As an example, TITIN (20.552 mutations) is a key component of striated muscles and mutations in this protein are related to different types of cardiomyopathies and muscular dystrophies (Hackman *et al*., 2002; Itoh-Satoh *et al*., 2002; Matsumoto *et al*., 2005). BRCA1, BRCA2 and APC (9.172, 12.063 and 9.237 mutations respectively) are proteins involved in DNA repair and tumor suppressor (Kawasaki *et al*., 2007; Liu *et al*., 2010; Shukla *et al*., 2011). It is well known mutations in these proteins produce an increased risk for different types of cancer, especially breast, ovarian and colorectal cancer (Easton *et al*., 2007; Mersch *et al*., 2015; Yamaguchi *et al*., 2016). Mutations do not appear equally in different protein regions, IDR and LCR have more mutations than the ordered portion of the protein (**Supplementary Figure 3**).

**Figure 1:**
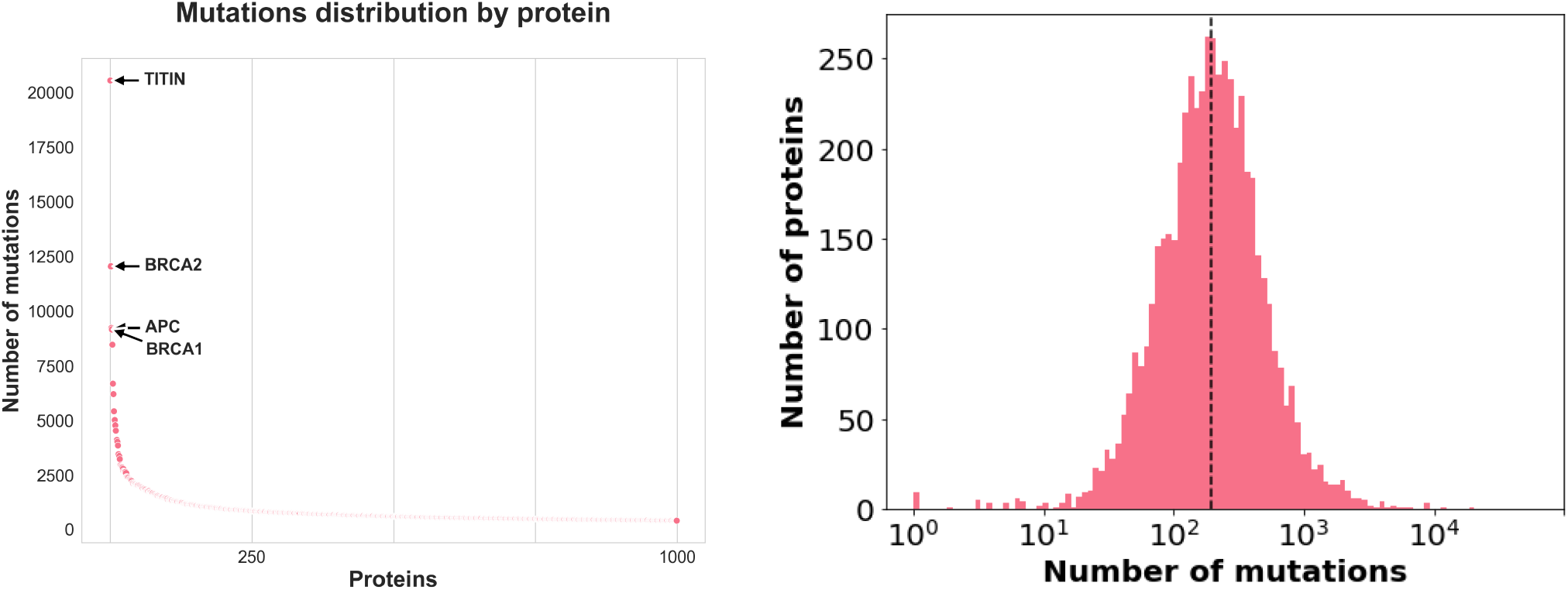
On the left, number of mutations by protein (only the first 1000 most mutated proteins are shown). On the right: Distribution of the number of mutations.

Each protein is associated with one or more LLPS source databases and, when possible, with their role in the LLPS process. Protein roles can vary depending on the MLO and the source database leading to diverse situations. A protein can be annotated as Driver in a particular MLO and as Client in another, also a protein can have a role in one database and be unassigned or have a different one in another for the same MLO. There are 285 proteins classified as Drivers, 357 regulators, 3.157 potential clients, and 4.105 have no role assigned in their source databases or MLO (**Supplementary Figure 4** shows the distribution of proteins by their role and, disaggregated by MLO).

Mutated proteins of DisPhaseDB are associated with 103 MLOs, varying in number across them. As an example, the nucleolus has 3.315 associated proteins while the synaptosome has only 1. Most proteins are associated with a single MLO (3.729), being the maximum 13 MLOs (1 protein) (**Figure 2**).

**Figure 2:**
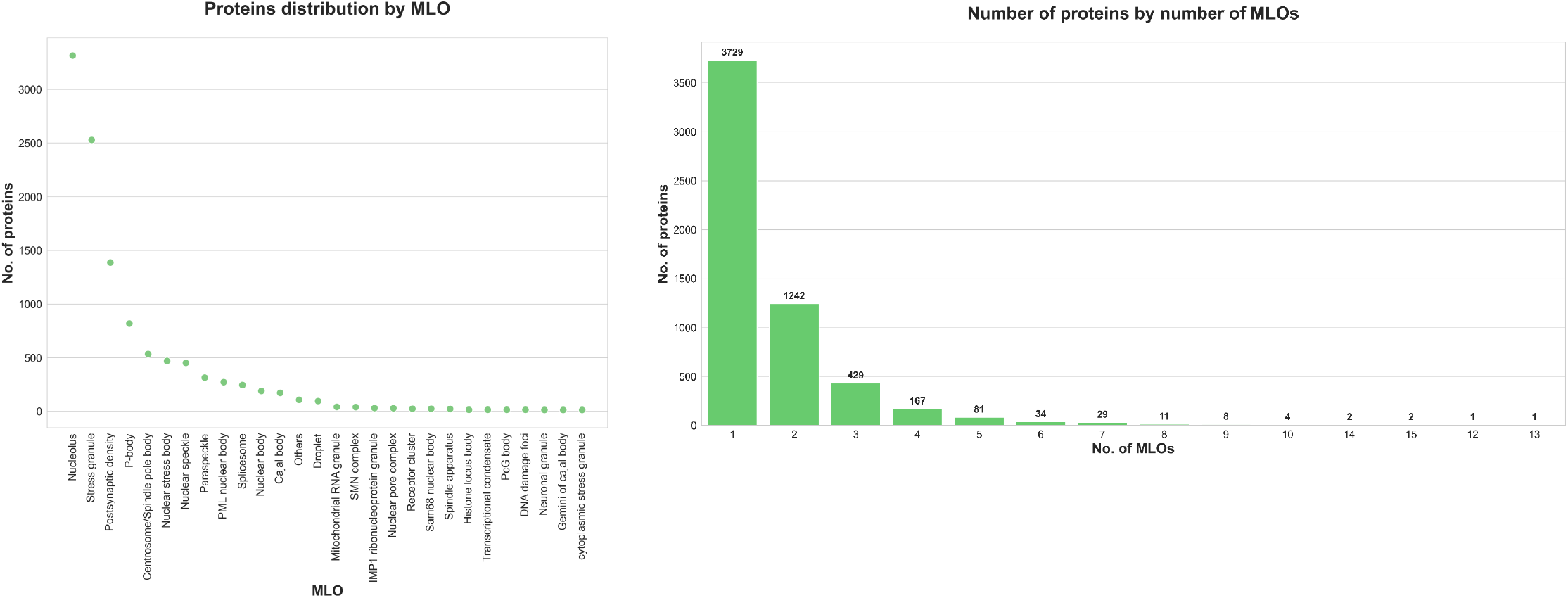
Left panel, protein distribution by MLO in DisPhaseDB, showing only MLOs with more than 10 associated proteins. Right panel: number of MLOs in which a protein can be present.

Also mutated proteins are associated with one or more diseases, **Figure 3** (upper panel) shows the number of DisPhaseDB mutated proteins associated with all the Mesh ontology subheadings in the disease category. These headings are nodes near the root of the ontology, but the annotations allow going forward to more specificity, for example **Supplementary Figure 5** shows the terms under “neoplasms” subheading disaggregated by site. Since 80% of the mutations in DisPhaseDB are contributed by COSMIC (somatic mutations in cancer). **Figure 3** (lower panel) shows the distribution of mutated proteins by disease removing those mutations contributed by COSMIC. Even though removing COSMIC mutations, proteins associated with neoplasms are still predominant.

**Figure 3:**
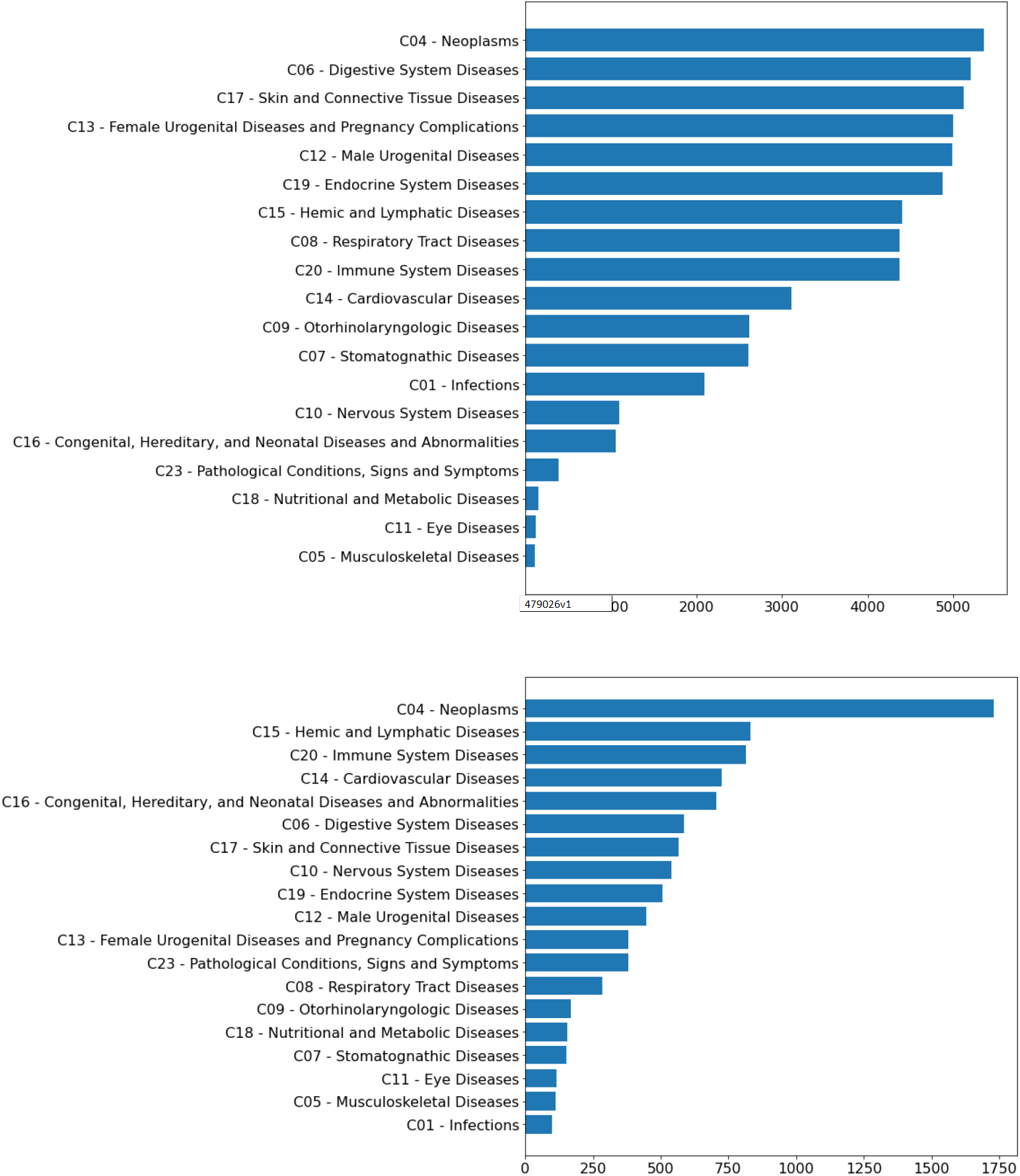
Upper panel, distribution of the total mutated proteins among all the subheading in MeSh ontology. Bottom panel, same as upper panel, but excluding COSMIC contributed mutations to see the tendencie (COSMIC contributes with 1.464.124 mutations out of 1.660.059)

### Server usage

**DisPhaseDB** offers either a quick search by protein, MLO or disease or an advanced search applying one or several filters. Possible fIlters are by protein, role, MLO, disease name or keyword, by evidence (low or high throughput experiments), by protein disorder content and mutation type (missense, frameshift, nonsense, etc). In addition, filters can be combined in such a way that users can customize the set of proteins according to their need or interest.

As an example, a protein search, Synaptic functional regulator FMR1 (UP: Q06787), displays the following outputs: section I) a protein summary with general information and fasta sequence; II) protein MLO location; III) protein features mapped onto the sequence such as regions, domains, disorder content and mutations (disaggregated by type), among others IV) mutation summary and V) a mutation table to download. **Figure 4** shows sections III and IV of the output as an example.

**Figure 4:**
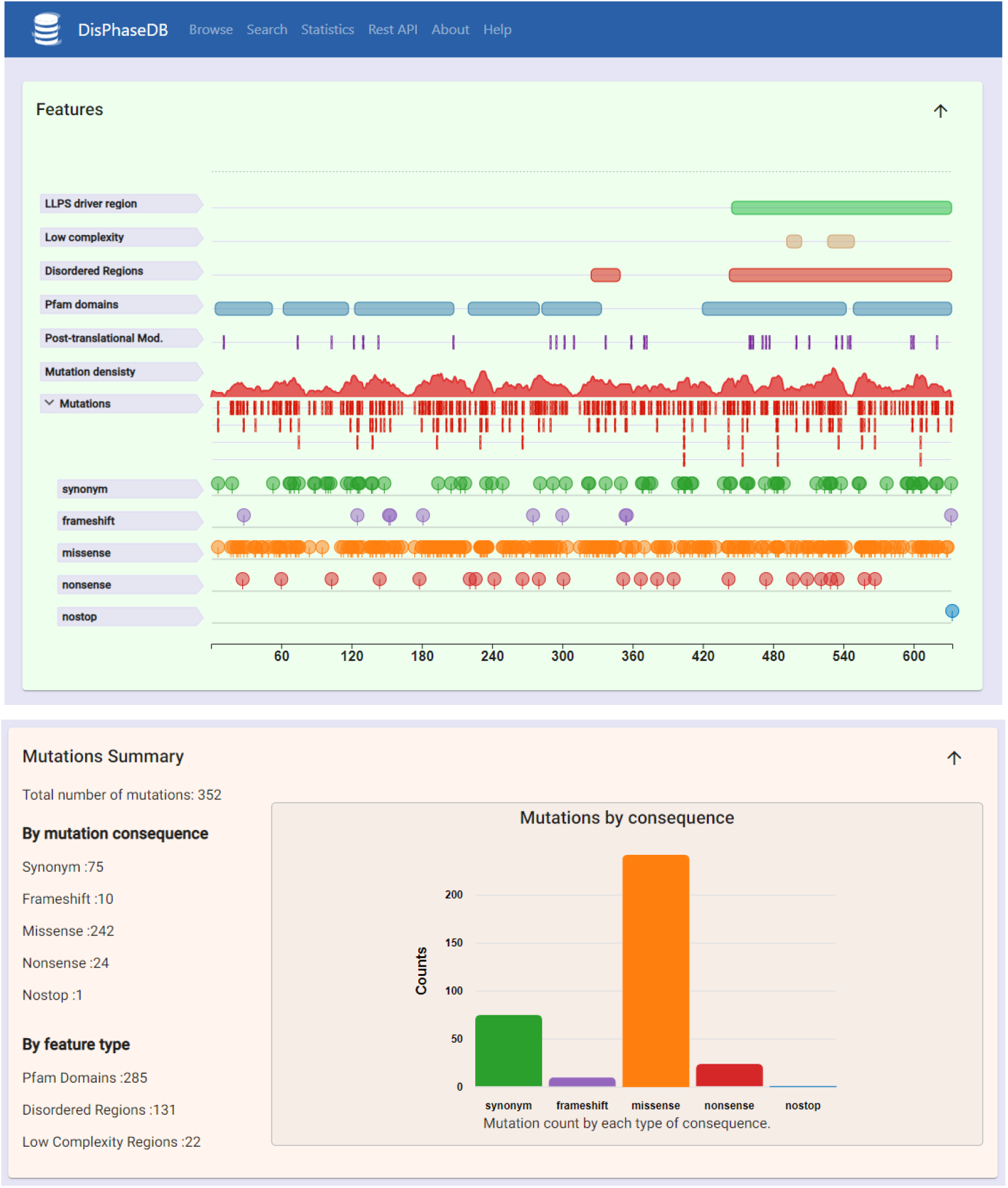
example of sections III and IV of the output for a search by protein, FMR1.

Other examples of searches can be retrieving all the proteins that are classified by role or in a particular disease. Another example, proteins associated with a particular MLO, the database will bring the proteins related to this MLO regardless of their role and disease.

## Discussion

To the best of our knowledge, there is not an integrated and comprehensive resource for mutations in MLOs associated proteins. For this reason, we integrated all state-of-the-art resources of proteins involved in LLPS and MLOs with four relevant disease databases that annotate medical terms and phenotypic effects. The variants’ databases selected are not redundant showing very little overlap among them. Furthermore, there are many mutation databases which makes it difficult to cover the range of diseases or effects with a single one. They are often not user friendly and they cross-reference to different ontologies and many other databases. This highlights the need for a unification of these resources.

It also provides mutation mapping over the sequence and metadata associated with the proteins, as disordered, low complexity and ordered regions, post translational modifications, among other features. Therefore this resource will be helpful to understand sequence-function relationships and mutational impact.

We expect DisPhaseDB to assist researchers to better understand complex human diseases under the lens of phase separation.

## Notes

https://disphasedb.leloir.org.ar

### Competing Interest Statement

The authors have declared no competing interest.

## References

Alberti, S. et al. (2009) ‘A systematic survey identifies prions and illuminates sequence features of prionogenic proteins’, Cell, 137(1), pp. 146–158.

Alberti, S. and Dormann, D. (2019) ‘Liquid-Liquid Phase Separation in Disease’, Annual review of genetics, 53, pp. 171–194.

Alderson, T.R. and Kay, L.E. (2021) ‘NMR spectroscopy captures the essential role of dynamics in regulating biomolecular function’, Cell, 184(3), pp. 577–595.

Amberger, J.S. et al. (2018) ‘OMIM.org: leveraging knowledge across phenotype–gene relationships’, Nucleic acids research, 47(D1), pp. D1038–D1043.

Banani, S.F. et al. (2017) ‘Biomolecular condensates: organizers of cellular biochemistry’, Nature reviews. Molecular cell biology, 18(5), pp. 285–298.

Boeynaems, S. et al. (2018) ‘Protein Phase Separation: A New Phase in Cell Biology’, Trends in cell biology, 28(6), pp. 420–435.

Brangwynne, C.P. et al. (2009) ‘Germline P granules are liquid droplets that localize by controlled dissolution/condensation’, Science, 324(5935), pp. 1729–1732.

Buniello, A. et al. (2019) ‘The NHGRI-EBI GWAS Catalog of published genome-wide association studies, targeted arrays and summary statistics 2019’, Nucleic acids research, 47(D1), pp. D1005–D1012.

Dignon, G.L., Best, R.B. and Mittal, J. (2020) ‘Biomolecular Phase Separation: From Molecular Driving Forces to Macroscopic Properties’, Annual review of physical chemistry, 71, pp. 53–75.

Easton, D.F. et al. (2007) ‘A systematic genetic assessment of 1,433 sequence variants of unknown clinical significance in the BRCA1 and BRCA2 breast cancer-predisposition genes’, American journal of human genetics, 81(5), pp. 873–883.

Gadd, J.C. et al. (2008) ‘Sizing subcellular organelles and nanoparticles con?ned within aqueous droplets’, Analytical chemistry, 80(9), pp. 3450–3457.

Gomes, E. and Shorter, J. (2019) ‘The molecular language of membraneless organelles’, The Journal of biological chemistry, 294(18), pp. 7115–7127.

Gopal, P.P. et al. (2017) ‘Amyotrophic lateral sclerosis-linked mutations increase the viscosity of liquid-like TDP-43 RNP granules in neurons’, Proceedings of the National Academy of Sciences of the United States of America, 114(12), pp. E2466–E2475.

Hackman, P. et al. (2002) ‘Tibial muscular dystrophy is a titinopathy caused by mutations in TTN, the gene encoding the giant skeletal-muscle protein titin’, American journal of human genetics, 71(3), pp. 492–500.

Harrison, A.F. and Shorter, J. (2017) ‘RNA-binding proteins with prion-like domains in health and disease’, Biochemical Journal, 474(8), pp. 1417–1438.

Hofweber, M. and Dormann, D. (2019) ‘Friend or foe—Post-translational modifications as regulators of phase separation and RNP granule dynamics’, The Journal of biological chemistry, 294(18), pp. 7137–7150.

Hornbeck, P.V. et al. (2019) ‘15 years of PhosphoSitePlus®: integrating post-translationally modified sites, disease variants and isoforms’, Nucleic acids research, 47(D1). doi:10.1093/nar/gky1159.

Itoh-Satoh, M. et al. (2002) ‘Titin mutations as the molecular basis for dilated cardiomyopathy’, Biochemical and biophysical research communications, 291(2), pp. 385–393.

Kawasaki, Y. et al. (2007) ‘Identification and characterization of Asef2, a guanine–nucleotide exchange factor specific for Rac1 and Cdc42’, Oncogene, pp. 7620–7627. doi:10.1038/sj.onc.1210574.

Köhler, S. et al. (2021) ‘The Human Phenotype Ontology in 2021’, Nucleic acids research, 49(D1), pp. D1207–D1217.

Landrum, M.J. et al. (2018) ‘ClinVar: improving access to variant interpretations and supporting evidence’, Nucleic acids research, 46(D1), pp. D1062–D1067.

van der Lee, R. et al. (2014) ‘Classification of intrinsically disordered regions and proteins’, Chemical reviews, 114(13), pp. 6589–6631.

Lenard, A.J. et al. (2021) ‘Phosphorylation Regulates CIRBP Arginine Methylation, Transportin-1 Binding and Liquid-Liquid Phase Separation’, Frontiers in molecular biosciences, 8, p. 689687.

Li, M.J. et al. (2012) ‘GWASdb: a database for human genetic variants identified by genome-wide association studies’, Nucleic acids research, 40(Database issue), pp. D1047–54.

Li, Q. et al. (2019) ‘LLPSDB: a database of proteins undergoing liquid–liquid phase separation in vitro’, Nucleic acids research, 48(D1), pp. D320–D327.

Liu, J. et al. (2010) ‘Human BRCA2 protein promotes RAD51 filament formation on RPA-covered single-stranded DNA’, Nature structural & molecular biology, 17(10), pp. 1260–1262.

Luo, R. et al. (2021) ‘A novel missense variant in ACAA1 contributes to early-onset Alzheimer’s disease, impairs lysosomal function, and facilitates amyloid-β pathology and cognitive decline’, Signal transduction and targeted therapy, 6(1), p. 325.

Luo, Y.-Y., Wu, J.-J. and Li, Y.-M. (2021) ‘Regulation of liquid-liquid phase separation with focus on post-translational modifications’, Chemical communications, 57(98), pp. 13275–13287.

Mao, Y.S., Zhang, B. and Spector, D.L. (2011) ‘Biogenesis and function of nuclear bodies’, Trends in genetics: TIG, 27(8), pp. 295–306.

Matsumoto, Y. et al. (2005) ‘Functional analysis of titin/connectin N2-B mutations found in cardiomyopathy’, Journal of muscle research and cell motility, 26(6-8), pp. 367–374.

Mersch, J. et al. (2015) ‘Cancers associated with BRCA1 and BRCA2 mutations other than breast and ovarian’, Cancer, 121(2), pp. 269–275.

Mészáros, B. et al. (2020) ‘PhaSePro: the database of proteins driving liquid-liquid phase separation’, Nucleic acids research, 48(D1), pp. D360–D367.

Mittag, T. and Parker, R. (2018) ‘Multiple Modes of Protein–Protein Interactions Promote RNP Granule Assembly’, Journal of molecular biology, 430(23), pp. 4636–4649.

Mungall, C.J. et al. (2017) ‘The Monarch Initiative: an integrative data and analytic platform connecting phenotypes to genotypes across species’, Nucleic acids research, 45(D1), pp. D712–D722.

Murakami, T. et al. (2015) ‘ALS/FTD Mutation-Induced Phase Transition of FUS Liquid Droplets and Reversible Hydrogels into Irreversible Hydrogels Impairs RNP Granule Function’, Neuron, 88(4), pp. 678–690.

Nelson, S.J. (2009) ‘Medical terminologies that work: The example of MeSH’, in 2009 10th International Symposium on Pervasive Systems, Algorithms, and Networks. 2009 10th International Symposium on Pervasive Systems, Algorithms, and Networks, IEEE. doi:10.1109/i-span.2009.84.

Ning, W. et al. (2020) ‘DrLLPS: a data resource of liquid-liquid phase separation in eukaryotes’, Nucleic acids research, 48(D1), pp. D288–D295.

Orti, F. et al. (2021) ‘Insight into membraneless organelles and their associated proteins: Drivers, Clients and Regulators’, Computational and structural biotechnology journal, 19, pp. 3964–3977.

Paladin, L. et al. (2020) ‘The Feature-Viewer: a visualization tool for positional annotations on a sequence’, Bioinformatics, 36(10), pp. 3244–3245.

Piñero, J. et al. (2020) ‘The DisGeNET knowledge platform for disease genomics: 2019 update’, Nucleic acids research, 48(D1), pp. D845–D855.

Piovesan, D. et al. (2021) ‘MobiDB: intrinsically disordered proteins in 2021’, Nucleic acids research, 49(D1), pp. D361–D367.

Prasad, A. et al. (2019) ‘Molecular Mechanisms of TDP-43 Misfolding and Pathology in Amyotrophic Lateral Sclerosis’, Frontiers in molecular neuroscience, 12, p. 25.

Ryan, V.H. and Fawzi, N.L. (2019) ‘Physiological, Pathological, and Targetable Membraneless Organelles in Neurons’, Trends in neurosciences, 42(10), pp. 693–708.

Sanders, D.W. et al. (2020) ‘Competing Protein-RNA Interaction Networks Control Multiphase Intracellular Organization’, Cell, 181(2), pp. 306–324.e28.

Schisa, J.A. and Elaswad, M.T. (2021) ‘An Emerging Role for Post-translational Modifications in Regulating RNP Condensates in the Germ Line’, Frontiers in molecular biosciences, 8, p. 658020.

Schmidt, H.B. and Görlich, D. (2015) ‘Nup98 FG domains from diverse species spontaneously phase-separate into particles with nuclear pore-like permselectivity’, eLife, 4. doi:10.7554/eLife.04251.

Schriml, L.M. et al. (2019) ‘Human Disease Ontology 2018 update: classification, content and work?ow expansion’, Nucleic acids research, 47(D1), pp. D955–D962.

Shin, Y. and Brangwynne, C.P. (2017) ‘Liquid phase condensation in cell physiology and disease’, Science, 357(6357). doi:10.1126/science.aaf4382.

Shukla, P.C. et al. (2011) ‘BRCA1 is an essential regulator of heart function and survival following myocardial infarction’, Nature communications, 2, p. 593.

Specht, C.G. (2021) ‘A Quantitative Perspective of Alpha-Synuclein Dynamics - Why Numbers Matter’, Frontiers in synaptic neuroscience, 13, p. 753462.

Su, Q., Mehta, S. and Zhang, J. (2021) ‘Liquid-liquid phase separation: Orchestrating cell signaling through time and space’, Molecular cell, 81(20), pp. 4137–4146.

Tang, S.-J. (2017) ‘Potential Role of Phase Separation of Repetitive DNA in Chromosomal Organization’, Genes, 8(10). doi:10.3390/genes8100279.

Tate, J.G. et al. (2019) ‘COSMIC: the Catalogue Of Somatic Mutations In Cancer’, Nucleic acids research, 47(D1), pp. D941–D947.

Tsang, B. et al. (2020) ‘Phase Separation as a Missing Mechanism for Interpretation of Disease Mutations’, Cell, 183(7), pp. 1742–1756.

UniProt Consortium (2021) ‘UniProt: the universal protein knowledgebase in 2021’, Nucleic acids research, 49(D1), pp. D480–D489.

Vernon, R.M. et al. (2018) ‘Pi-Pi contacts are an overlooked protein feature relevant to phase separation’, eLife, 7. doi:10.7554/eLife.31486.

Wu, X. et al. (2020) ‘Liquid-Liquid Phase Separation in Neuronal Development and Synaptic Signaling’, Developmental cell, 55(1), pp. 18–29.

Yamaguchi, K. et al. (2016) ‘Reduced expression of APC-1B but not APC-1A by the deletion of promoter 1B is responsible for familial adenomatous polyposis’, Scientific Reports. doi:10.1038/srep26011.

You, K. et al. (2020) ‘PhaSepDB: a database of liquid-liquid phase separation related proteins’, Nucleic acids research, 48(D1), pp. D354–D359.

