## supplementary information for "DisPhaseDB, an integrative database of diseases related variations in liquid-liquid phase separation proteins"

### Supplementary material

##### Supplementary Figure 1: Mutations among disease variant databases.

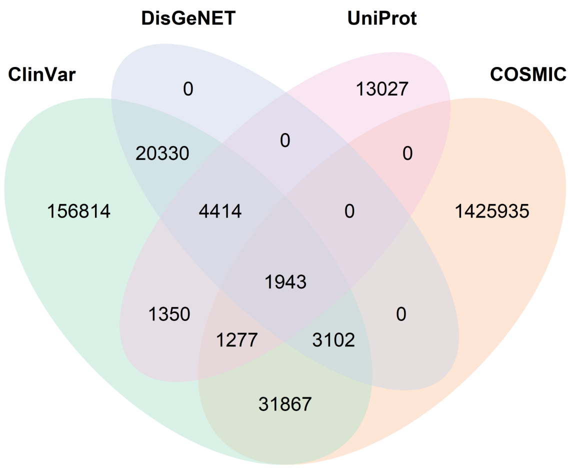


**Suppl. Fig. 1:** Overlap of coding mutations among disease dedicated databases. Only 1.943 out of 1.660.059 mutations are shared between the 4 databases.

##### Supplementary Figure 2. Proportion of mutations by type in DisPhaseDB

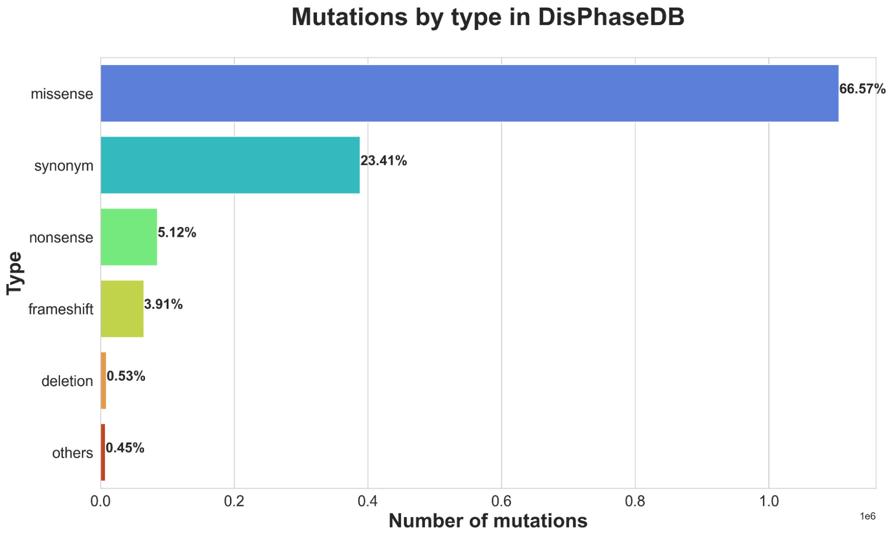

**Suppl. Figure 2:** Number of mutations by type in DisPhaseDB.

There are 10 possible types of mutations in DisPhaseDB proteins based on HGVS recommendations: missense, synonym, nonsense, frameshift, deletions, and in a lesser extent delins, insertions, duplications, no-stop and repeated (grouped as "others" for simplicity).

#### Supplementary Figure 3 proportion of mutations by region

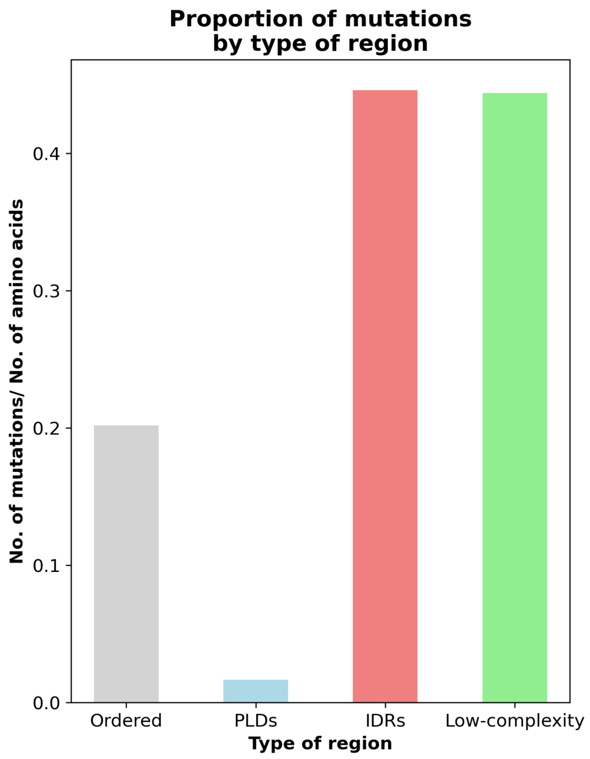


**Suppl. Figure 3:** Proportion of mutations by region (number of mutations/total number of residues in this type of region).

#### Supplementary Figure 4: Roles of the proteins in DisPhaseDB

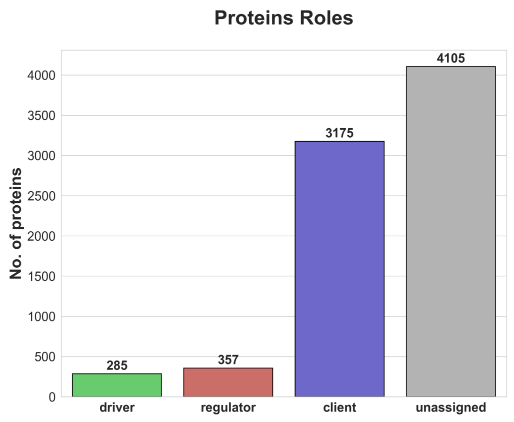


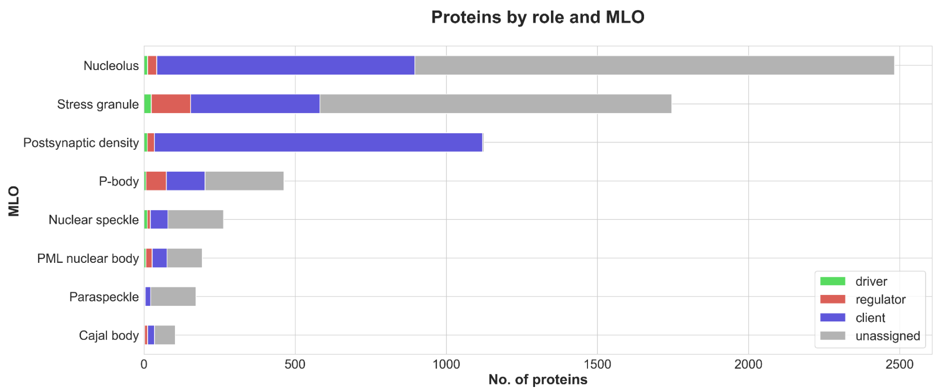


**Suppl. Figure 4:** Upper panel: proteins by their role. bottom panel: proteins by their role disagregated by MLOs.

**Supplementary Figure 5: Frequency of mutation by proteins associated with a disease term.**

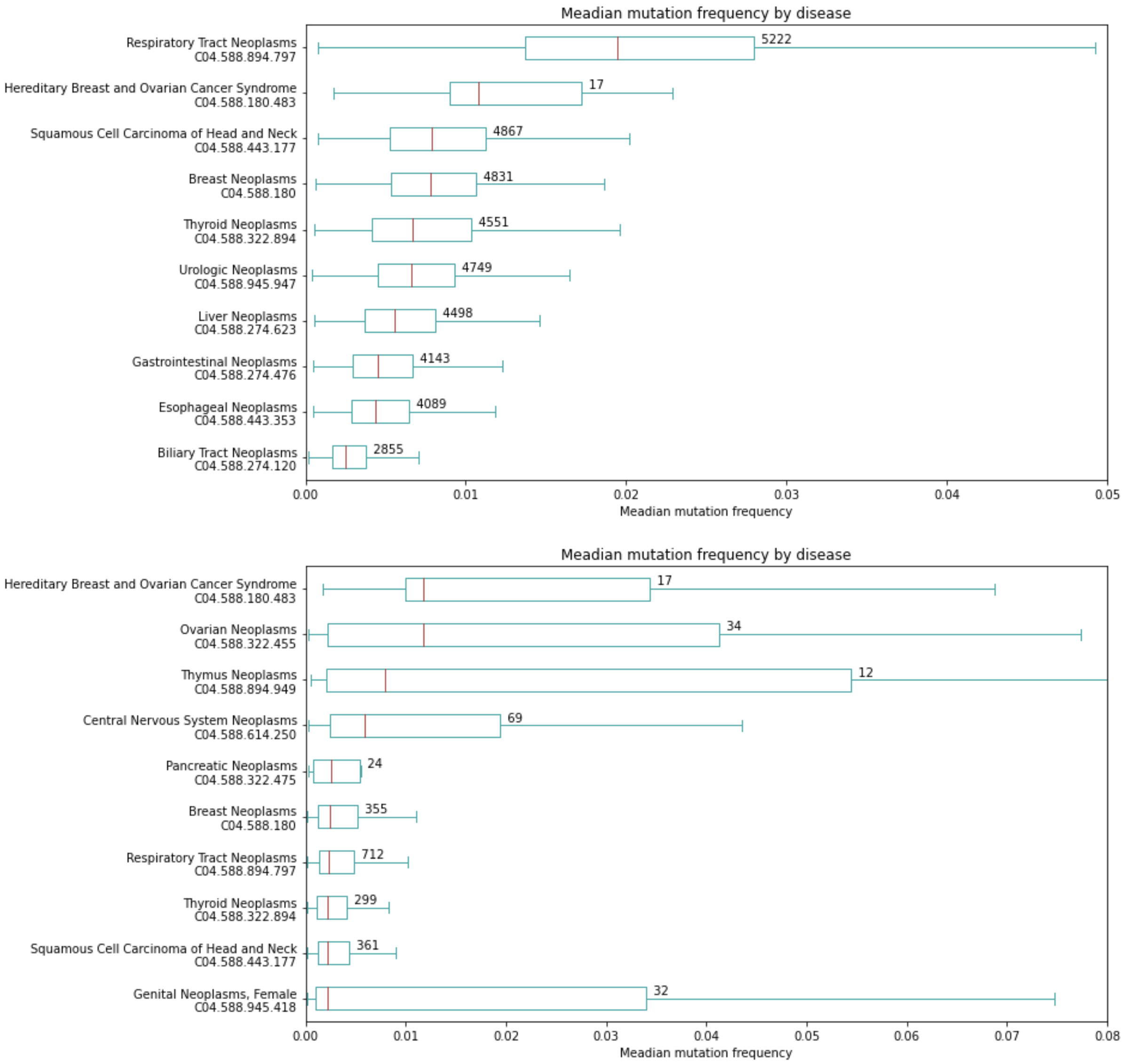


**Suppl. Figure 5: Mutation frequency in neoplasms by site MeSH subgraph**. We mapped all terms to childs of the subgraph of MeSH corresponding to "neoplasms by site" (tree number C04.588) with a depth of maximum 4 steps from the root. For each mapped term we collect the proteins having missense mutations associated with this term from DisPhaseDB. The boxplot shows the distribution of mutation frequency for each disease term (number of mutations divided by protein length). The number at the right of the boxes indicates the number of proteins in the set. Upper panel: all the mutations; lower panel, excluding COSMIC mutations.
